# Genomic evidence for the degradation of terrestrial organic matter by pelagic Arctic Ocean Chloroflexi bacteria

**DOI:** 10.1101/325027

**Authors:** David Colatriano, Patricia Tran, Celine Guéguen, Williams J. William, Connie Lovejoy, David A. Walsh

## Abstract

The Arctic Ocean currently receives a large supply of global river discharge and terrestrial dissolved organic matter. Moreover, an increase in freshwater runoff and riverine transport of organic matter to the Arctic Ocean is a predicted consequence of thawing permafrost and increased precipitation. The fate of the terrestrial humic-rich organic material and its impact on the marine carbon cycle are largely unknown. Here, the first metagenomic survey of the Canada Basin in the Western Arctic Ocean showed that pelagic Chloroflexi from the Arctic Ocean are replete with aromatic compound degradation genes, acquired in part by lateral transfer from terrestrial bacteria. Our results imply marine Chloroflexi have the capacity to use terrestrial organic matter and that their role in the carbon cycle may increase with the changing hydrological cycle.

## Introduction

The Arctic Ocean accounts for 1.4% of global ocean volume but receives 11 % of global river discharge ^1^. Up to 33 % of the dissolved organic matter in the Arctic Ocean is of terrestrial origin ^2^ and a major fraction of this terrestrial dissolved organic matter (tDOM) originates from carbon-rich soils and peatlands ^3,4^. With thawing permafrost and increased precipitation occurring across the Arctic ^5^, increases in freshwater runoff and riverine transport of organic matter to the Arctic Ocean are predicted, which will increase tDOM fluxes and loadings ^6,7^. The additional tDOM may represent new carbon and energy sources for the Arctic Ocean microbial community and contribute to increased respiration, which would result in the Arctic being a source of dissolved inorganic carbon to the ocean. Alternatively, as it moves from its source of origin to the Arctic Ocean tDOM could become more recalcitrant to bacterial metabolism and represent a long term sequestration of the newly released carbon making the Arctic more carbon neutral ^8,9^. However, an estimated 50% of Arctic Ocean tDOM is removed before being released to the Atlantic, at least in part by microbial processes ^10^. As input of tDOM increases, knowledge on its microbial transformation will be critical for understanding changes in Arctic carbon cycling.

The marine SAR202 is a diverse and uncultivated clade of Chloroflexi bacteria that comprise roughly 10% of planktonic cells in the dark ocean ^11–15^. SAR202 is also common in marine sediments and deep lakes ^16–18^. It has long been speculated that SAR202 may play a role in the degradation of recalcitrant organic matter ^12,15^, and the recent analysis of SAR202 single-cell amplified genomes (SAGs) lends support to this notion^19^. More generally, Chloroflexi, including those in the SAR202 clade, are also present in the upper layers of the Arctic Ocean ^20^, leading to the hypothesis that recalcitrant organic compounds present in high Arctic tDOM could be utilized by this group.

## Results

In this study, we analyzed Chloroflexi metagenome assembled genomes (MAGs) generated from samples collected from the vertically stratified waters of the Canada Basin in the Western Arctic Ocean (Fig. 1a). A metagenomic co-assembly was generated from samples originating from the surface layer (5 m to 7 m), the subsurface chlorophyll maximum (25 m to 79 m) and a layer corresponding to the terrestrially-derived DOM fluorescence (FDOM) maximum previously described within the cold CB halocline comprised of Pacific-origin waters (177 m to 213 m) ^21^. The Pacific-origin FDOM maximum is due to sea ice formation and interactions with bottom sediments on the Beaufort and Chukchi shelves, which themselves are influenced by coastal erosion and river runoff ^21^. Binning based on tetranucleotide frequency and coverage resulted in 360 MAGs from a diversity of marine microbes (Fig. 1b). Six near complete Chloroflexi MAGs were identified. Based on 16S rRNA gene phylogeny, these MAGs represented 3 distinct SAR202 subclades (SAR202-II, -VI, -VII), the AncK29 clade and the TK10 clade (Fig. 2a). Estimated MAG completeness ranged from 77 % to 99 %, while contamination ranged from 0 % to 2.3 % (Table 1). All MAGs exhibited highest coverage just below the subsurface chlorophyll maximum (Fig. 2b) which is consistent with earlier findings on SAR202 distribution in the North Pacific Ocean ^13^. However, the concentration and composition of the FDOM maximum in the CB is significantly different compared to the North Pacific Ocean ^22^ and the North Atlantic Subtropical Gyre ^23^. A concatenated protein phylogeny demonstrated that the SAR202 MAGs were distinct from previously published MAGs from the deep ocean ^19^ and oxygen minimum zones ^24^ (Supplementary Fig. 1). Fragment recruitment of 21 TARA Ocean metagenomic datasets spanning epipelagic to mesopelagic waters at 7 locations and 4 separate bathypelagic metagenomes indicated that the CB Chloroflexi MAGs were not widely distributed in the oceans (Fig. 2c, Supplementary Data 1). These findings are evidence that the Chloroflexi MAGs represent genotypes that are rare outside Arctic marine waters.

**Table 1.** Genomic characteristics of MAGs.

|  | Size<br>(Mb) | Cov (x) | GC (%) | Completeness<br>(%) | Contamination<br>(%) | N50<br>(kb) | # of<br>Contigs |
| --- | --- | --- | --- | --- | --- | --- | --- |
| **SAR202-II-3** | 1.36 | 68 | 39 | 80 | 0 | 35 | 46 |
| **SAR202-II-177A** | 1.62 | 101 | 42 | 82 | 1 | 36 | 52 |
| **SAR202-VII-2** | 2.78 | 16 | 59 | 99 | 0 | 246 | 8 |
| **SAR202-VI-29A** | 1.52 | 16 | 46 | 97 | 2 | 59 | 2 |
| **TK10-74A** | 2.45 | 12 | 69 | 81 | 2.3 | 38 | 76 |
| **Anck29-46** | 1.11 | 11 | 32 | 77 | 0 | 75 | 22 |

**Fig. 1.**
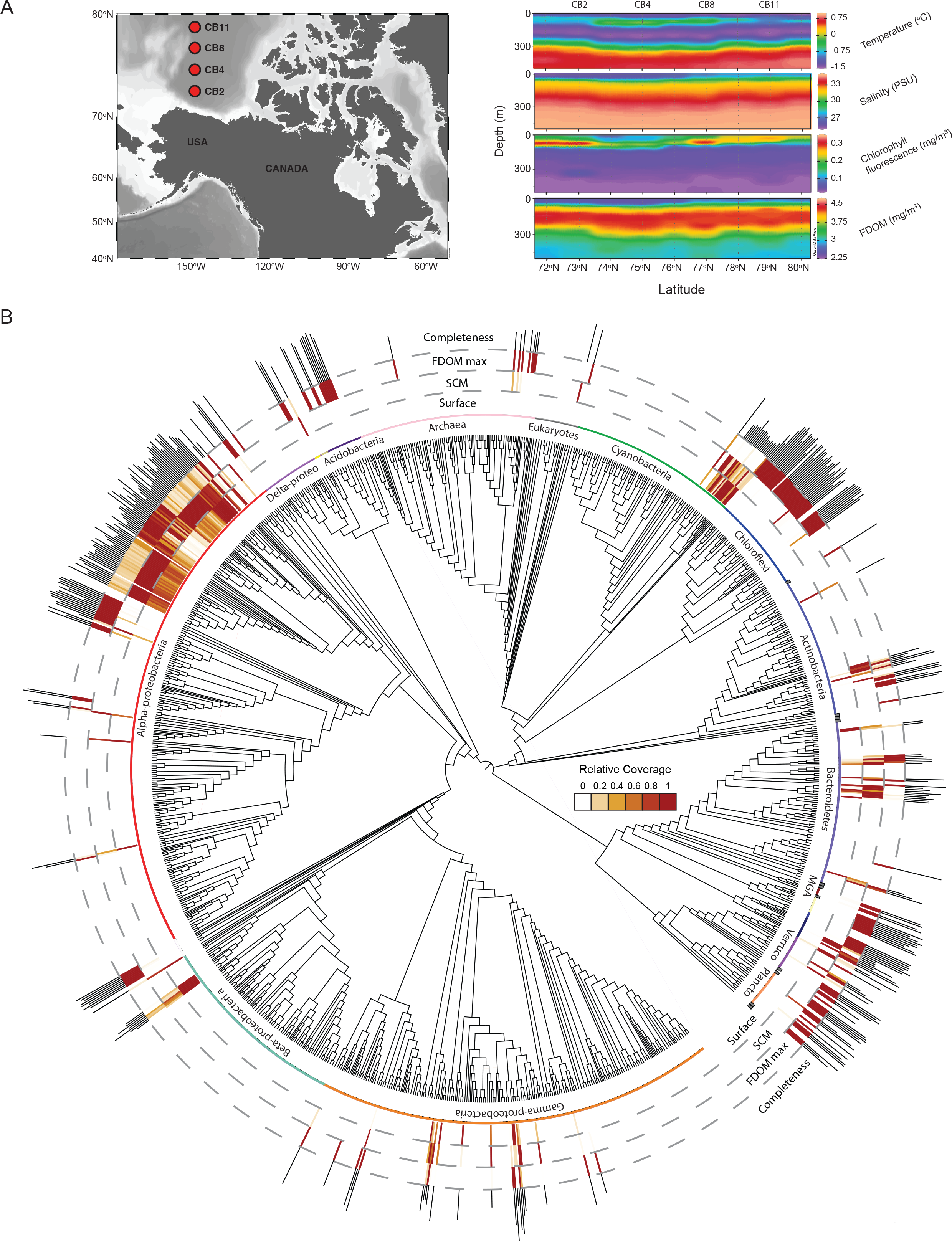
Metagenomic survey of microbial diversity in the Canada Basin. (**A**) Sampling locations and environmental profiles of the Canada Basin generated with Ocean Data Viewer. (**B**) Concatenated protein phylogeny of 360 Arctic Ocean MAGs inferred by MetaWatt and visualized in iTOL. The three inner tracks present relative coverage of MAGs averaged across samples collected from surface waters, subsurface chlorophyll maximum (SCM) and the fluorescent dissolved organic matter maximum (FDOM max). The outer track presents estimated MAG completeness as inferred by MetaWatt. MAG completeness ranged from 25-94%.

**Fig. 2.**
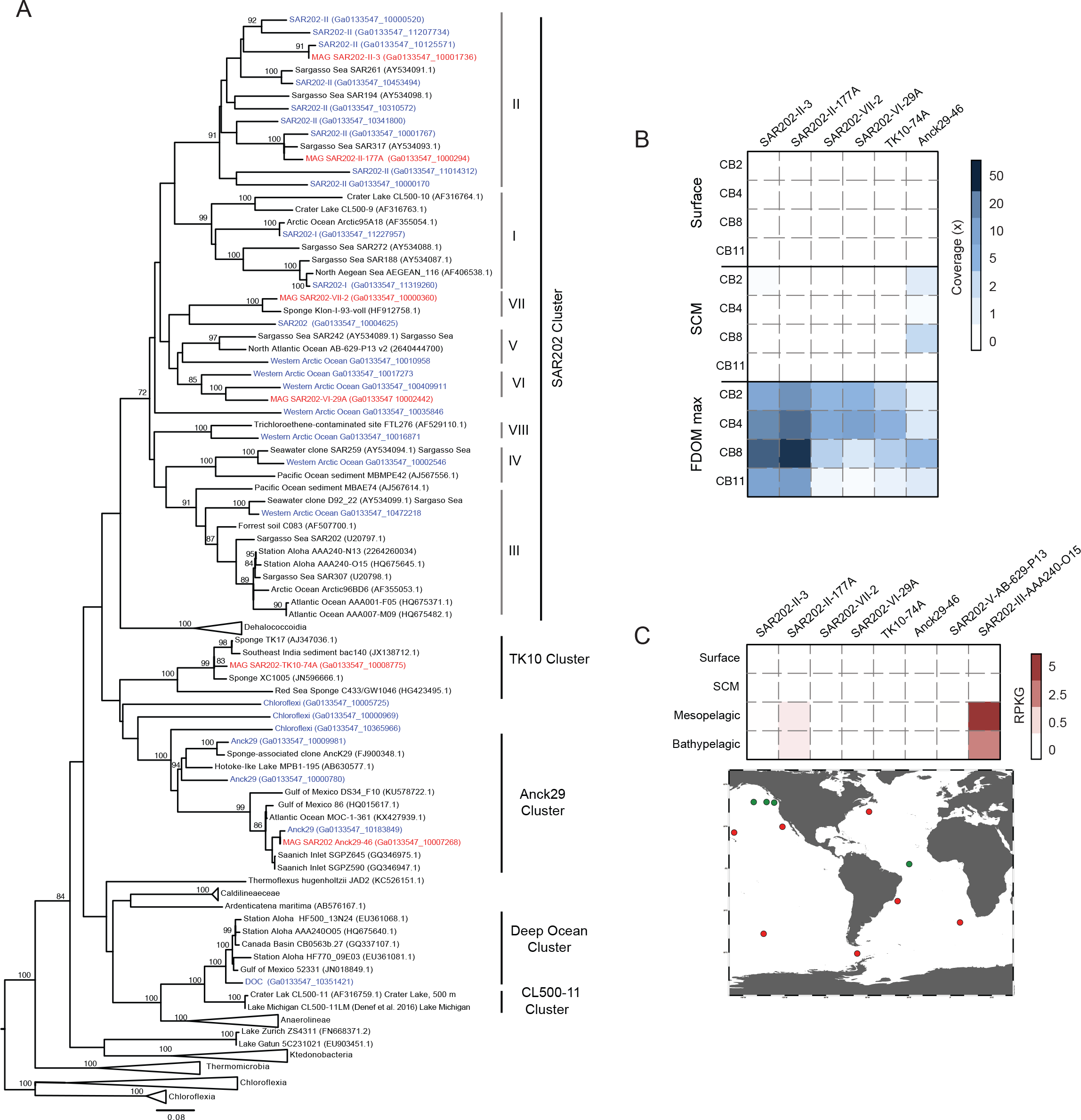
Diversity and distribution of Arctic Ocean Chloroflexi MAGs. (**A**) Maximum likelihood tree of Chloroflexi based on partial 16S rRNA gene sequences. Blue taxa are from Canada Basin metagenomes and red taxa are from the six Chloroflexi MAGs. Bootstrap values of >70% are shown (100 replicates) (**B**) Distribution of MAGs in the Canada Basin based on metagenome coverage at the surface, subsurface chlorophyll maximum (SCM) and fluorescent dissolved organic matter maximum (FDOM max) (**C**) Distribution of MAGs in global ocean metagenomes based on fragment recruitment. Two deep ocean SAR202 SAGs from Landry et al. ^11^ were included for comparison. Location of TARA ocean metagenomes (red) and bathypelagic metagenomes (green) are shown on the map.

The Chloroflexi MAGs contained many genes implicated in the degradation of aromatic compounds typically associated with humic-rich tDOM (Supplementary Data 2). A single MAG (SAR202-VII-2) from a previously undescribed clade (SAR202-VII) exhibited a striking enrichment in these genes (Fig. 3a). Partial pathways for the catabolism of aromatic compounds were recently reported from deep ocean SAR202 SAGs ^19^. To assess whether the abundance and diversity of SAR202-VII-2 genes involved in aromatic compound catabolism is unique to Arctic Ocean MAGs or is a more broad characteristic of marine Chloroflexi, we compared gene content between SAR202-VII-2 and two SAGs (SAR202-V-AB-629-P13 and SAR202-III-AAA240-O15) reported in ^19^. Of the 117 SAR202-VII-2 orthologs implicated in aromatic compound degradation, 12 were identified in SAR202-III-AAA240-O15 and only one was identified in SAR202-V-AB-629-P13, implying distinct and less diverse pathways in deep ocean SAR202 compared to the Arctic Ocean populations (Supplementary Data 2).

**Fig. 3.**
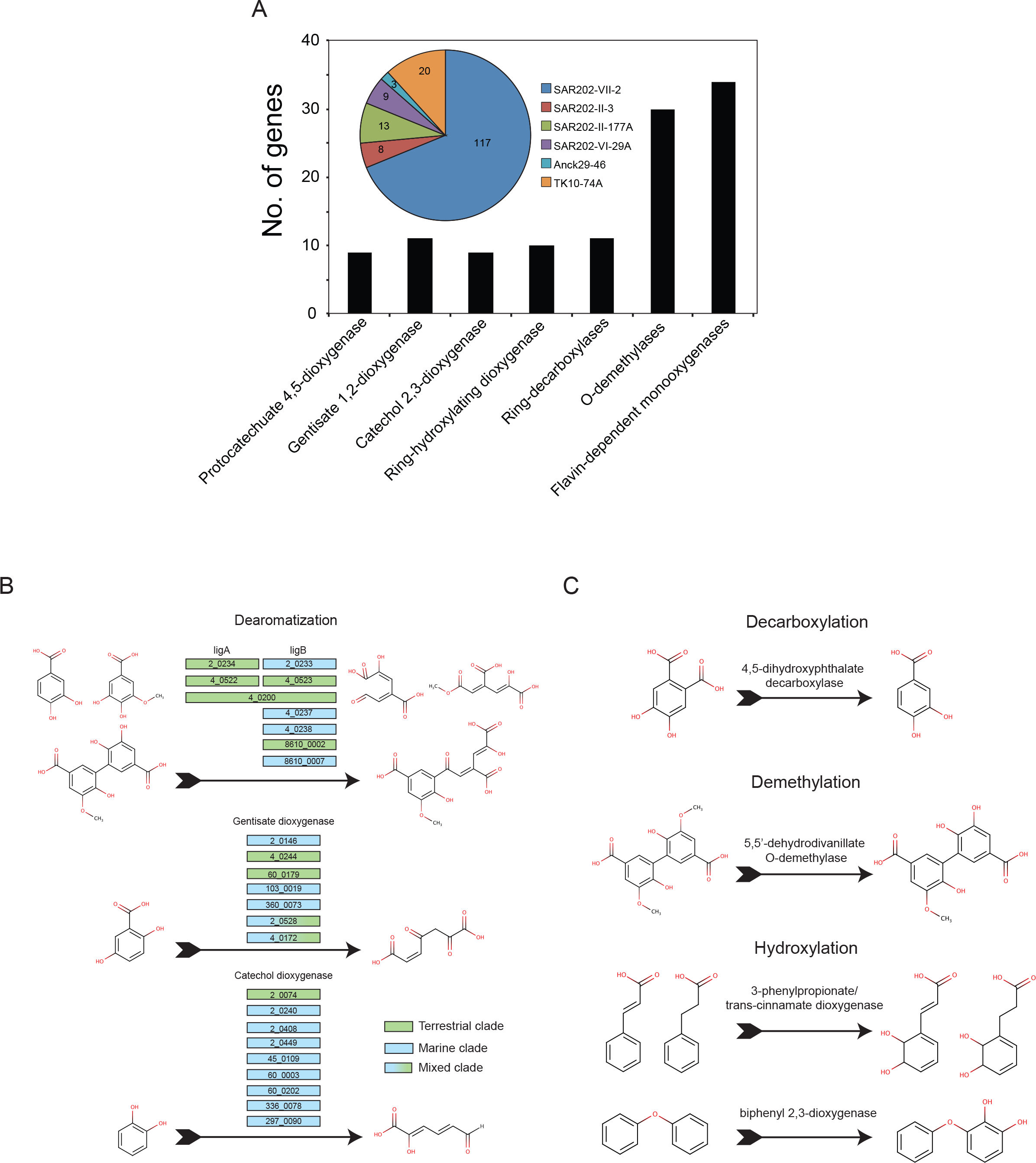
Aromatic compound degradation genes and pathways in Arctic Ocean Chloroflexi MAGs. (**A**) Abundance of aromatic compound degradation genes in Chloroflexi MAGs from the Canada Basin (pie chart) and a breakdown of those found specifically in SAR202-VII-2 (column plot) (**B**) Predicted aromatic ring-opening enzymatic reactions identified in SAR202-VII-2 with gene loci displayed in the coloured boxes. Genes in green and blue boxes were most closely related to homologs from terrestrial or marine bacteria, respectively. Genes in the blue/green boxes were in clades containing diverse environmental bacteria. (**C**) Examples of predicted aromatic ring-modifying enzymatic reactions identified in SAR202-VII-2.

Proteins for the modification and degradation of monoaryl and biaryl compounds were predicted, including a diversity of aromatic ring-cleaving dioxygenases ^25–27^. A total of 42 ring-cleaving dioxygenases targeting compounds related to catechol, protocatechuate and gentisate were present in the six MAGs, with 25 dioxygenases predicted in SAR202-VII-2 alone (Fig. 3a-b). Ring demethylation, hydroxylation and decarboxylation are important prerequisite steps to prime diverse aromatic compounds for downstream oxidative cleavage ^28,29^. Thirty ring-demethylating monooxygenases, ten ring-hydroxylating dioxygenases, and eleven ring-decarboxylases were annotated in the SAR202-VII-2 MAG (Fig. 3c, Supplementary Data 2). Proteins involved in the conversion of ring-cleavage products to central intermediates of the citric acid cycle were also present in the SAR202-VII-2 MAG, including dehydrogenases (*i.e.* 2,3-dihydroxy-2,3-dihydrophenylpropionate dehydrogenase), decarboxylases (*i.e.* oxaloacetate B-decarboxylase), aldolases (*i.e.* HMG aldolase and 4-carboxymuconolactone decarboxylase), hydratases (*i.e.* 4-oxalmescanoate hydratase and 2-oxopent-4-enoate hydratase), isomerases (*i.e.* mycothiol maleulpyruvate isomerase and muconolactone isomerase) and hydrolases (*i.e.* 3-oxoadipate enol-lactonase and β-ketoadipate enol-lactone hydrolase) (Supplementary Data 2, Supplementary Fig. 2). We note that we were unable to identify a single complete reference pathway for humic-like aromatic compound degradation. Since estimated genome completeness for SAR202-VII-2 was 99%, it is unlikely the genes were missed due to an incomplete genome. Another explanation is that marine Chloroflexi genomes encode novel pathway variants. Indeed, numerous metal-dependent hydrolases, hydrolases of the HAD family and NAD(P)-dependent dehydrogenase were clustered in genomic regions with the ring-modifying oxygenases, decarboxylases, and demethylases described above. In addition to the array of aromatic compound degradation genes, the SAR202-VII-2 MAG also contained 34 copies of the flavin mononucleotide (FM)/F420-dependent monooxygenase catalytic subunit (FMNO) proteins previously implicated in activation of recalcitrant organic compounds in the deep ocean ^19^. These results are consistent with Chloroflexi in the Arctic Ocean having the metabolic potential to access carbon and energy available in aromatic compounds typically associated with tDOM.

The diversity of SAR202-VII-2 genes implicated in aromatic compound degradation lead us to hypothesize that they may have originated by lateral gene transfer from terrestrial bacteria. To test this, we targeted aromatic compound degradation genes (the ring-cleaving dioxygenases, specifically) in the Chloroflexi MAGs for in-depth phylogenetic analyses. The genomic diversity of marine Chloroflexi was expanded in our analysis by including 130 Chloroflexi MAGs recently assembled and binned from the TARA Oceans project ^30^. A number of the SAR202-VII-2 ring-cleaving dioxygenase homologs were most closely related to proteins from the TARA Ocean Chloroflexi MAGs and other marine originating genomes, particularly the catechol dioxygenases (Supplementary Fig. 3), 3 gentisate 1,2 dioxygenases (Fig. 4) and methylgallate dioxygenases (Supplementary Fig. 4) indicating that aromatic compound degradation in Chloroflexi is not restricted to the Arctic Ocean. However, lateral gene transfer from terrestrial bacteria was also evident. For example, an annotated gentisate dioxygenase gene was positioned within a clade of terrestrial Actinomycetes (Fig. 4). Additional genes involved in the degradation of structures related to catechol, protocatechuate and gentisate were phylogenetically associated with homologs from terrestrial Acidobacteria (Supplementary Fig. 5), Actinobacteria (Supplementary Fig. 6), Armatimonadetes (Fig. 4), Delta-proteobacteria (Supplementary Fig. 3 and 6), Beta-proteobacteria (Supplementary Fig. 7) and a clade of diverse terrestrial phyla (Supplementary Fig. 8). Additionally, 2 gentisate 1,2-dioxygenase genes and 1 protocatechuate dioxygenase *ligB* gene were phylogenetically associated to a clade of genes from both terrestrial Delta-proteobacteria and marine microbes (Fig. 4 and Supplementary Fig. 8). These putative gene acquisitions were unlikely due to contaminating scaffolds because the genes were located on long scaffolds that were assigned to Chloroflexi with high confidence based on tetranucleotide frequencies and the phylogenetic identity of house-keeping genes. Such a phylogenetic pattern supports the hypothesis that marine Chloroflexi acquired the capacity for aromatic compound degradation, at least in part, by lateral gene transfer from terrestrial bacteria.

**Fig. 4.**
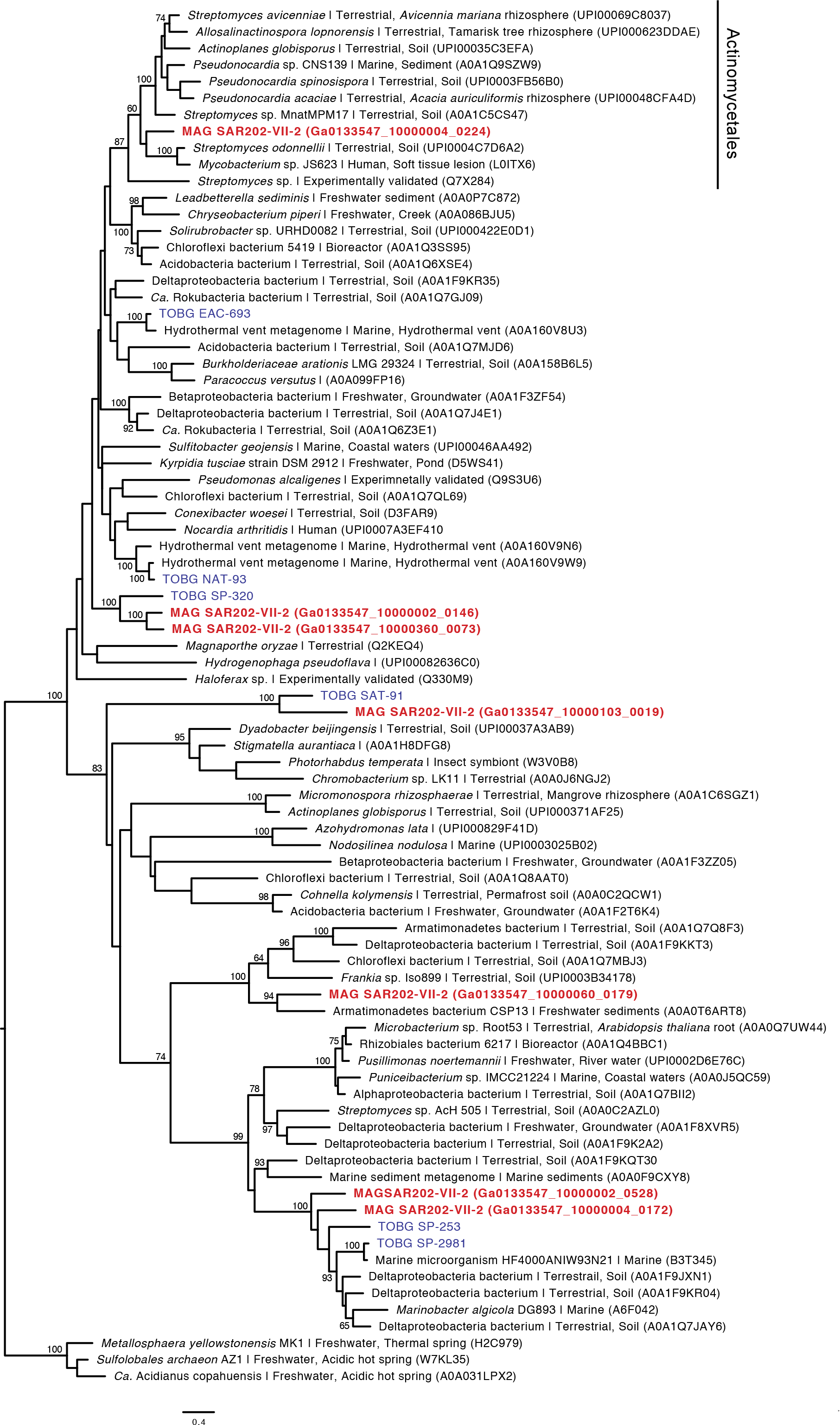
Maximum likelihood tree of predicted gentisate 1,2-dioxygenases. Maximum likelihood tree of predicted gentisate 1,2-dioxygenases. Bootstrap values of >60% are shown (100 replicates). Homologs from SAR202-VII-2 are highlighted in red and homologs from TARA Ocean MAGs are highlighted in blue.

## Conclusion

In total, these results are consistent with Chloroflexi playing a role in tDOM transformation in waters of the Arctic Ocean. This is the first study to our knowledge to associate a specific microbial group with tDOM metabolism in the Arctic Ocean and it expands on recent studies contributing to our understanding of the metabolic diversity of the abundant yet uncultivated marine Chloroflexi ^19,24^. Moreover, lateral gene transfer from terrestrial bacteria appears to have contributed to the evolution of aromatic compound degradation capabilities within marine Chloroflexi, particularly in regions of the Arctic Ocean impacted by terrestrial input.

The majority of MAGs were restricted to the humic-rich Pacific-origin halocline of the CB, however it is the surface waters that will be most immediately affected by increased freshwater input ^1^. Hence, our initial observations suggest a need for further research on the distribution of tDOM-utilizing microbes in other Arctic water masses with an aim to establish how common and phylogenetically widespread tDOM metabolism is in the Arctic Ocean. These water masses could include coastal surface waters at the mouth of the Mackenzie River, as well as regions of differing DOM composition such as the East Siberian Sea ^21^. Moreover, metagenomic studies such as this are, in essence, hypothesis-generating and future work that includes targeted cultivation, *in situ* gene expression analysis, and rate measurement-based approaches are required to validate and quantify microbial metabolic contributions to nutrient cycling. Overall it is likely that marine Chloroflexi have the capacity to degrade tDOM, and their role in the Arctic carbon cycle may increase as Arctic warming leads to greater inputs of terrestrial organic matter.

## Methods

### Sampling and DNA extraction

Twelve samples for metagenomics were collected in September 2015 during the Joint Ocean Ice Study cruise to the Canada Basin. For each sample, 4-8 L of seawater was sequentially filtered through a 50 μm pore mesh, followed by a 3 μm pore size polycarbonate filter and a 0.22 μm pore size Sterivex filter (Durapore; Millipore, Billerica, MA, USA). Filters were preserved in RNAlater and stored at −80 °C until processed in the laboratory. DNA was extracted from the Sterivex filter using the following method: filters were thawed on ice and RNAlater was removed. The Sterivex was then rinsed twice with a sucrose-based lysis buffer, and filled with 1.8 mL of the lysis buffer. Filters were treated with 100 μL of 125 mg mL^−1^ lysozyme and 20 μL of 10 μg mL^−1^ RNAse A and left to rotate at 37 °C for 1 hour. After incubation, 100 μL of 10 mg mL^−1^ proteinase K and 100 μL of 20% SDS was added. Filters were left to rotate for 2 hours at 55 °C. Lysate was removed from the filters. Protein was precipitated and removed with 0.583 volumes of MPC Protein Precipitation Reagent (Epicentre, Madison, WI, USA) and centrifugation at 10,000 g at 4 °C for 10 minutes. The supernatant was transferred to a clean tube. DNA was precipitated with cold isopropanol, and resuspended in low TE buffer.

### Metagenomic sequencing, assembly, annotation, and binning

DNA sequencing of 12 samples was performed at the Department of Energy Joint Genome Institute (Walnut Creek, CA, USA) on the HiSeq 2500-1TB (Illumina) platform. Paired-end sequences of 2 × 150bp were generated for all libraries. A metagenome coassembly of all raw reads was generated using MEGAHIT ^31^ with kmer sizes of 23,43,63,83,103,123. Gene prediction and annotation was performed using the DOE Joint Genome Institute’s Integrated Microbial Genomes (IMG) database tool (version 4.11.0) ^32^. Metagenomic binning was performed on scaffolds greater than or equal to 10 Kb in length using MetaWatt ^33^. Relative weight of coverage binning was set to 0.75 and the optimize bins and polish bins options were set to on. The taxonomic identity of MAGs was assessed using a concatenated phylogenetic tree based on 138 single copy conserved genes as implemented in MetaWatt ^33^. Visualization of the tree and mapping of data on to taxa was performed with iTOL ^34^. Estimation of MAG completeness and contamination was performed using CheckM ^35^. Six Chloroflexi MAGs were selected for further analysis based on the presence of a 16S rRNA gene, high completeness and low contamination. Manual curation of the six Chloroflexi MAGs was performed and suspected contaminating scaffolds (single copy genes most similar to non-Chloroflexi taxa) were removed prior to further analysis of MAGs.

### 16S rRNA phylogenetic analysis

Chloroflexi diversity in the metagenomic assembly was assessed by 16S rRNA gene analysis. All 16S rRNA genes in the co-assembly were assigned to taxonomic groups using mothur ^36^ and the Wang method with a bootstrap value cutoff of 60 % ^37^. Chloroflexi 16S rRNA genes greater than 360 bp were included in a phylogenetic analysis with Chloroflexi reference sequences. A multiple sequence alignment was generated using MUSCLE (implemented in MEGA6) ^38^. Phylogenetic reconstructions were conducted by maximum likelihood using MEGA6-v.0.6 and the following settings: general time reversible model, gamma distribution model for the rate variation with four discrete gamma categories, and the nearest-neighbour interchange (NNI) heuristic search method ^39^ with a bootstrap analysis using 100 replicates.

### Single protein and concatenated protein phylogenies

A concatenated protein phylogeny was constructed using 30 Chloroflexi reference genomes and the 6 Canada Basin Chloroflexi MAGs. Orthologous genes in the 36 Chloroflexi genomes were identified using ProteinOrtho ^40^. Fifty orthologs present in at least 34 of the 36 genomes were selected for concatenated phylogenetic analysis (Supplemetary Data 3). Each orthologous protein family was aligned using MUSCLE (implemented in MEGA6) and alignment positions were masked using the probabilistic masker ZORRO ^41^, masking columns with weights less than 0.5. The concatenated alignment consisted of 14,815 amino acid positions. Phylogenetic reconstructions were conducted by maximum likelihood using MEGA6-v.0.6 and the following settings: JTT substitution model, gamma distribution with invariant sites model for the rate variation with four discrete gamma categories, and the nearest-neighbour interchange (NNI) heuristic search method ^39^ with a bootstrap analysis using 100 replicates.

For phylogenetic analysis of ring-cleaving dioxygenase sequences identified in SAR202-VII-2, query sequences were searched against UniRef90 and 130 Chloroflexi MAGs constructed from the TARA Oceans dataset ^30^. The TARA Ocean Chloroflexi MAGs were used as is with no manual curation. UniRef90 sequences and the top TARA Ocean MAG hits for each dioxygenase were aligned with their respective SAR202-VII-2 homologs with MUSCLE (implemented in MEGA6) and alignment positions were masked using the probabilistic masker ZORRO ^41^, masking columns with weights less than 0.5. Phylogenetic reconstruction was conducted using the same settings as the concatenated phylogeny.

### Comparative genomics and metabolic reconstruction

The distribution of orthologs across Arctic Ocean genomes, as well as the identification of orthologs shared with the deep ocean SAGs, was determined using proteinortho ^40^. Inference of protein function and metabolic reconstruction was based on the IMG annotations provided by the JGI, including KEGG, Pfam, EC numbers, and Metacyc annotations. Metabolic reconstruction was also facilitated by generated pathway genome databases for each MAG using the pathologic software available through Pathway Tools ^42^.

### Metagenomic fragment recruitment

The distribution of the Canada Basin MAGs in the global ocean was determined using best-hit reciprocal blast analysis similar to Landry et. al 2017 ^19^. Unassembled metagenomic data from 25 samples (Supplementary Data 1) was first recruited to the six Canada Basin Chloroflexi MAGS as well as two SAR202 SAGs originating from the deep North Pacific Ocean from the Hawaiian ocean time series (HOTS) and the deep North Atlantic Ocean ^19^. Metagenomes from the TARA Ocean project used here were representative of the surface, chlorophyll maximum and mesopelagic waters from the North Atlantic, South Atlantic, North Pacific, Coastal North Pacific, South Pacific, Coast of Brazil and the Antarctic peninsula. To reduce computational demand, only part 1 (1 Gbp of a random subset of reads) of each metagenomic dataset available at EBI was used (Supplementary Data 1). Additional bathypelagic metagenomes from the North Pacific and South Atlantic Oceans (LineP P04, P12 and P26, and Knorr S15 2500 m) were also included. All hits from the initial blast were then reciprocally queried against the Canada Basin Chloroflexi MAGs, bathypelagic SAR202 SAGs, and 130 Chloroflexi MAGs constructed from the TARA oceans data. The best hit was reported. Only hits with an alignment length greater than or equal to 100 bp and a percent identity of 95% or more were counted (lower % identity cut-offs did not alter the number of reads recruited in any significant manner). To compare the results among the different data sets, the number of recruited reads was normalized to total number of reads in each sample. The final coverage results were expressed as the number of reads per kilobase of the MAG per gigabase of metagenome (rpkg).

### Data Availability

The metagenomic data generated in this study are available in the Integrated Microbial Genomes database at the Joint Genome Institute at https://img.jgi.doe.gov, GOLD Project ID: Ga0133547. Metagenome assembled genomes are available at NCBI under the Bioproject XXXXXXXXX and accession numbers XXXXXX (for SAR202-II-3), XXXXXX (for SAR202-II-177A), XXXXXXX (for SAR202-VI-29A), XXXXXXX (for SAR202-VII-2), XXXXXX (for Anck29-46) and XXXXXX (for TK10-74A).

## Acknowledgments

The data were collected aboard the CCGS Louis S. St-Laurent in collaboration with researchers from Fisheries and Oceans Canada at the Institute of Ocean Sciences and Woods Hole Oceanographic Institution’s Beaufort Gyre Exploration Program and are available at http://www.whoi.edu/beaufortgyre. We would like to thank both the Captain and crew of the CCGS Louis S. St-Laurent and the scientific teams aboard. The work was conducted in collaboration with the U.S. Department of Energy Joint Genome Institute, a DOE Office of Science User Facility, and was supported under Contract No. DE-AC02-05CH11231. Funding from the Canadian Natural Science and Engineering Research Council (NSERC) Discovery grants (D.W., C.G. and C.L.) and the Canada Research Chair Program (D.W., C.G.) are acknowledged. D.C. was supported by FRQNT and Concordia’s Institute for Water, Energy and Sustainable Systems.

## Authors contributions

D.C. and D.W. designed and carried out the metagenomic experiments. W. J. W., C. L. and C. G. designed the sampling strategy. W. J. W. provided oceanographic data. D.W. collected samples. D.C. extracted the environmental genomic DNA and analyzed sequencing data. P. T. contributed to the bioinformatic analyses. D.C. and D.W. with contributions from C. G. and C. L. wrote the manuscript. All authors commented on the manuscript.

---

**Supplementary Fig 1.**
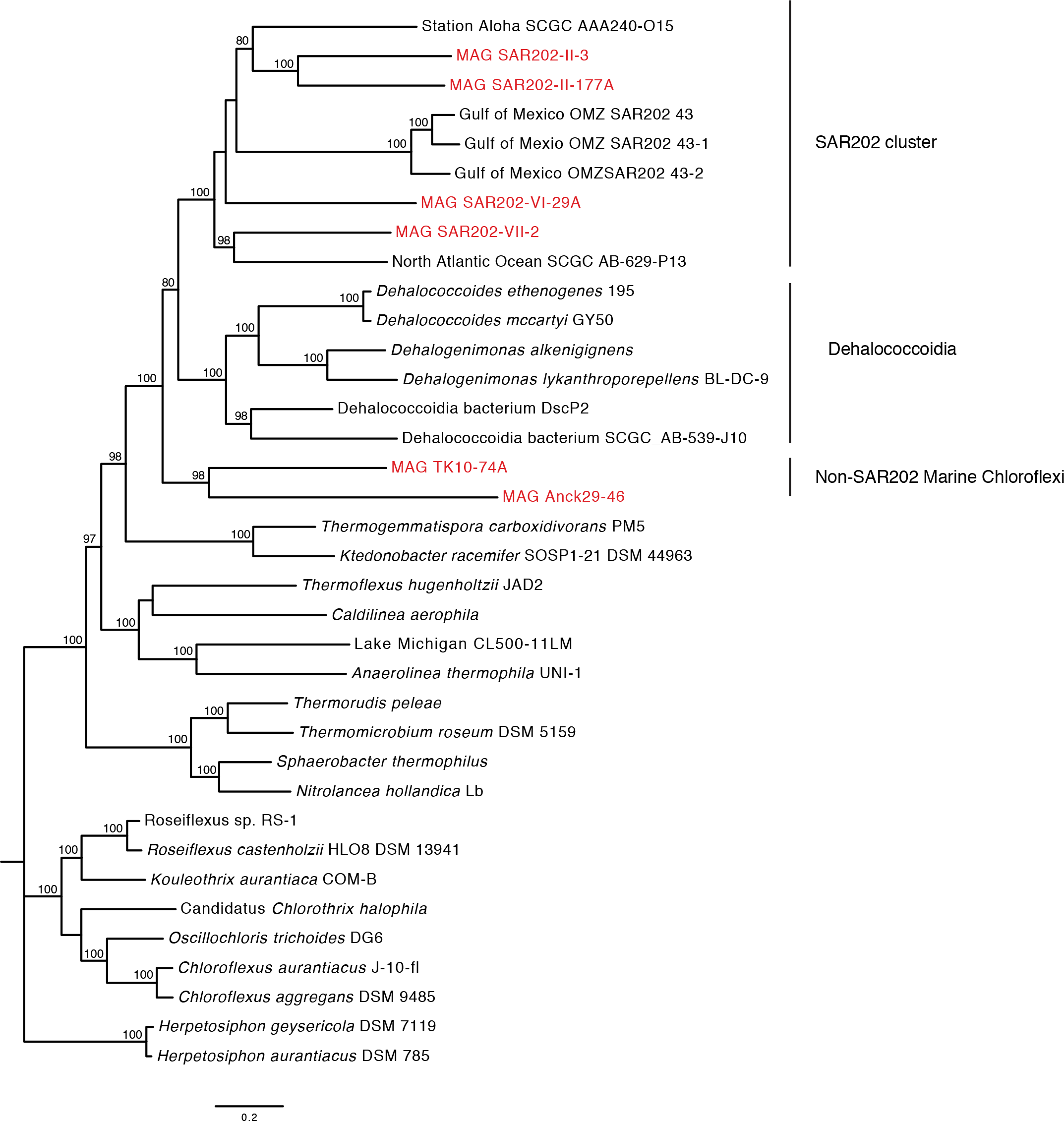
A concatenated protein phylogeny of Chloroflexi genomes including 30 Chloroflexi reference genomes and the 6 Canada Basin Chloroflexi MAGs. This is a maximum likelihood tree based on concatenated sequence alignment of fifty amino acid sequences (representing of 14,815 amino acid positions), present in at least 34 of the 36 Chloroflexi genomes. Bootstrap values are based on 100 replicates.

**Supplementary Fig 2.**
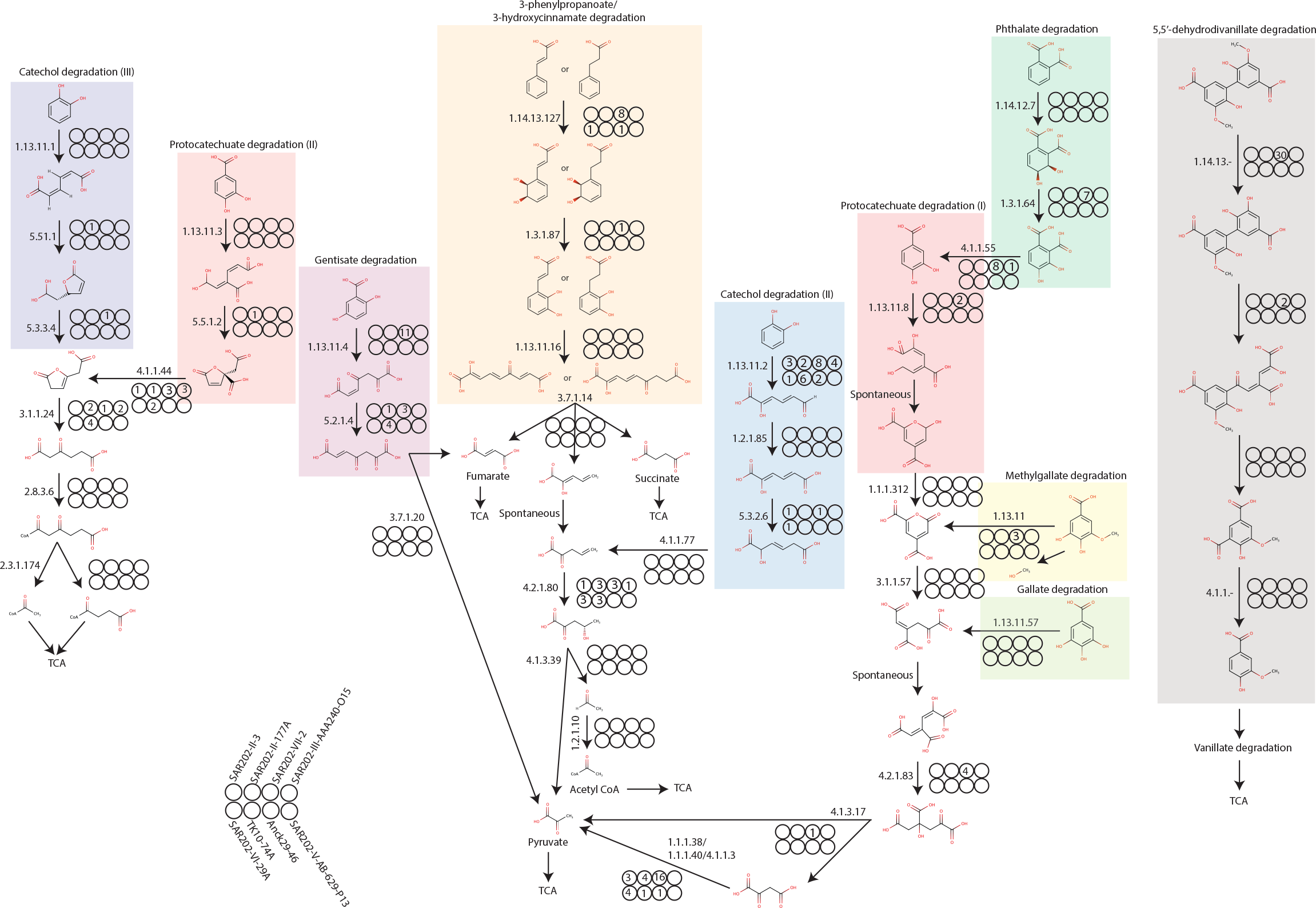
Aromatic compound degradation pathways identified in the Arctic Ocean and deep ocean Chloroflexi genomes. Circles represent the six Arctic Ocean MAGs and two deep ocean SAGs. The values in the circles represent the number of homologs identified in each respective genome. These reference metabolic pathways from MetaCyc were populated with proteins based on functional annotations available at the Integrated Microbial Genomes database.

**Supplementary Fig 3.**
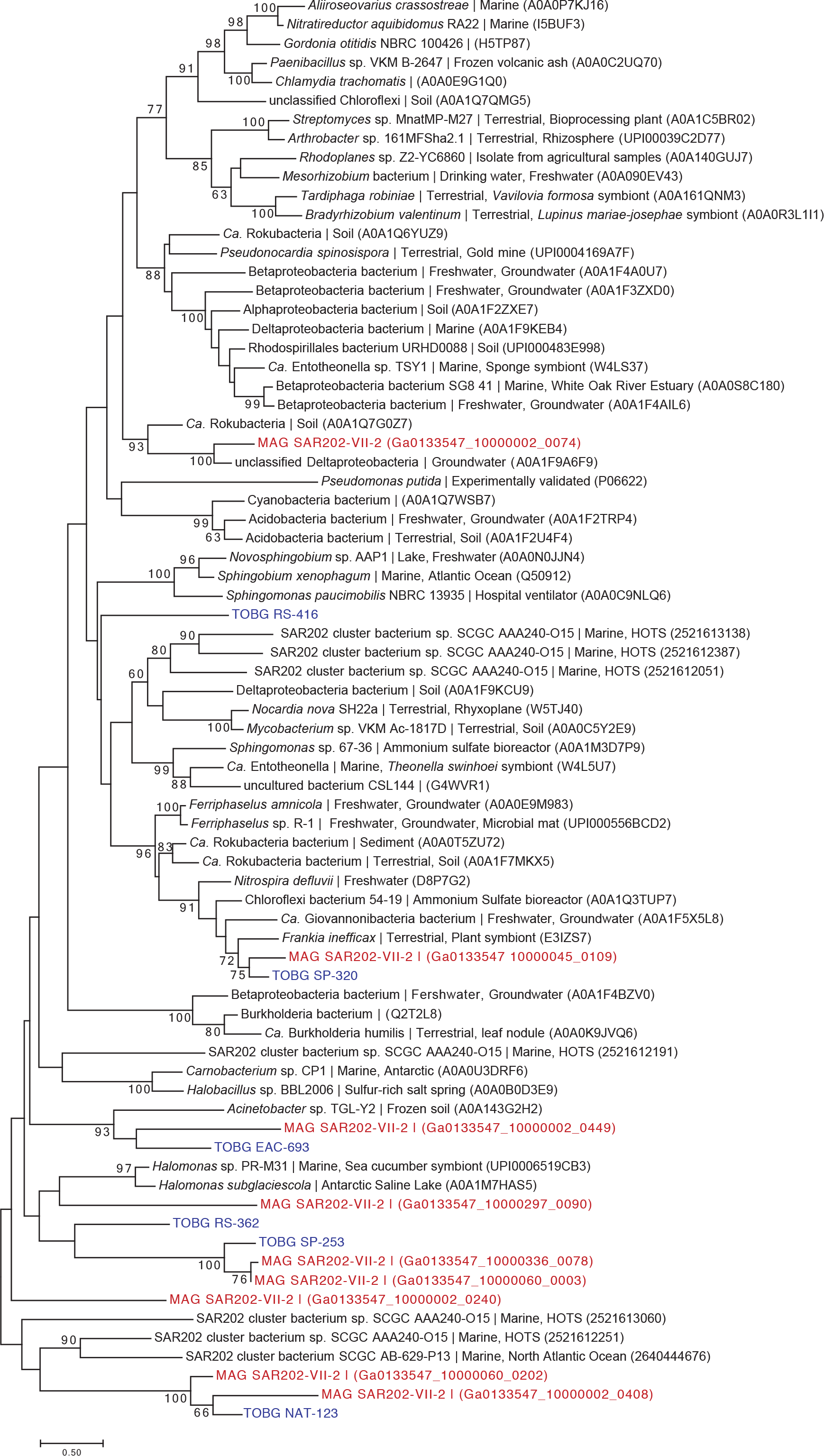
Phylogenetic analysis of predicted catechol 2,3-dioxygenase (pfam 00903) protein sequences identified in SAR202-VII-2. This is a maximum likelihood tree based the alignment of catechol 2,3-dioxygenase sequences identified in SAR202-VII-2, UniRef90 sequences and Chloroflexi MAGs constructed from the TARA Oceans dataset. Bootstrap values of >60% are shown (100 replicates). SAR202-VII-2 sequences are shown in red while sequences originating from the TARA Oceans MAGs are shown in blue

**Supplementary Fig 4.**
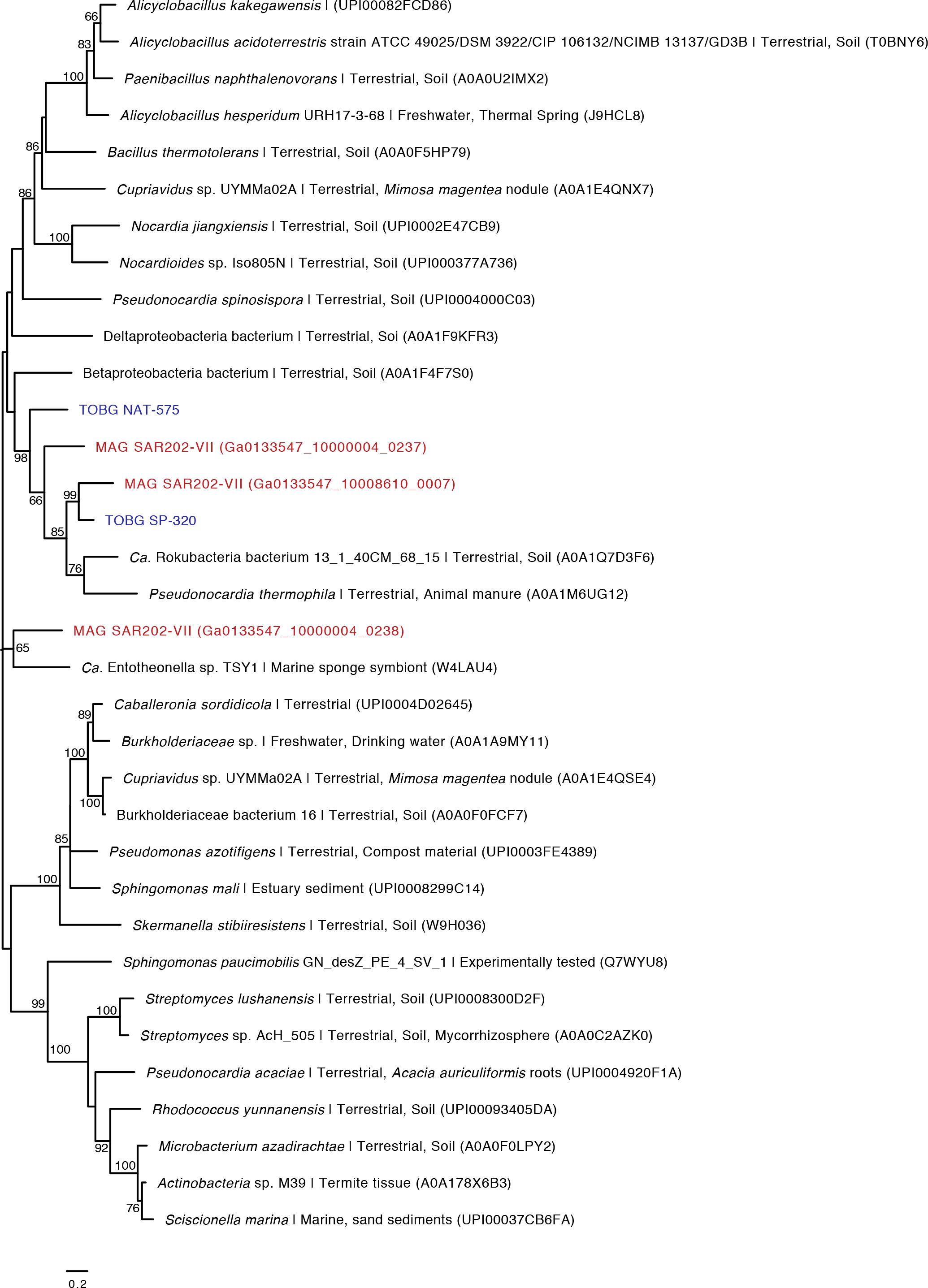
Phylogenetic analysis of predicted 3-O-methylgallate dioxygenase pfam 02900) protein sequences identified in SAR202-VII-2. Method and description as described in Supp Figure 2.

**Supplementary Fig 5.**
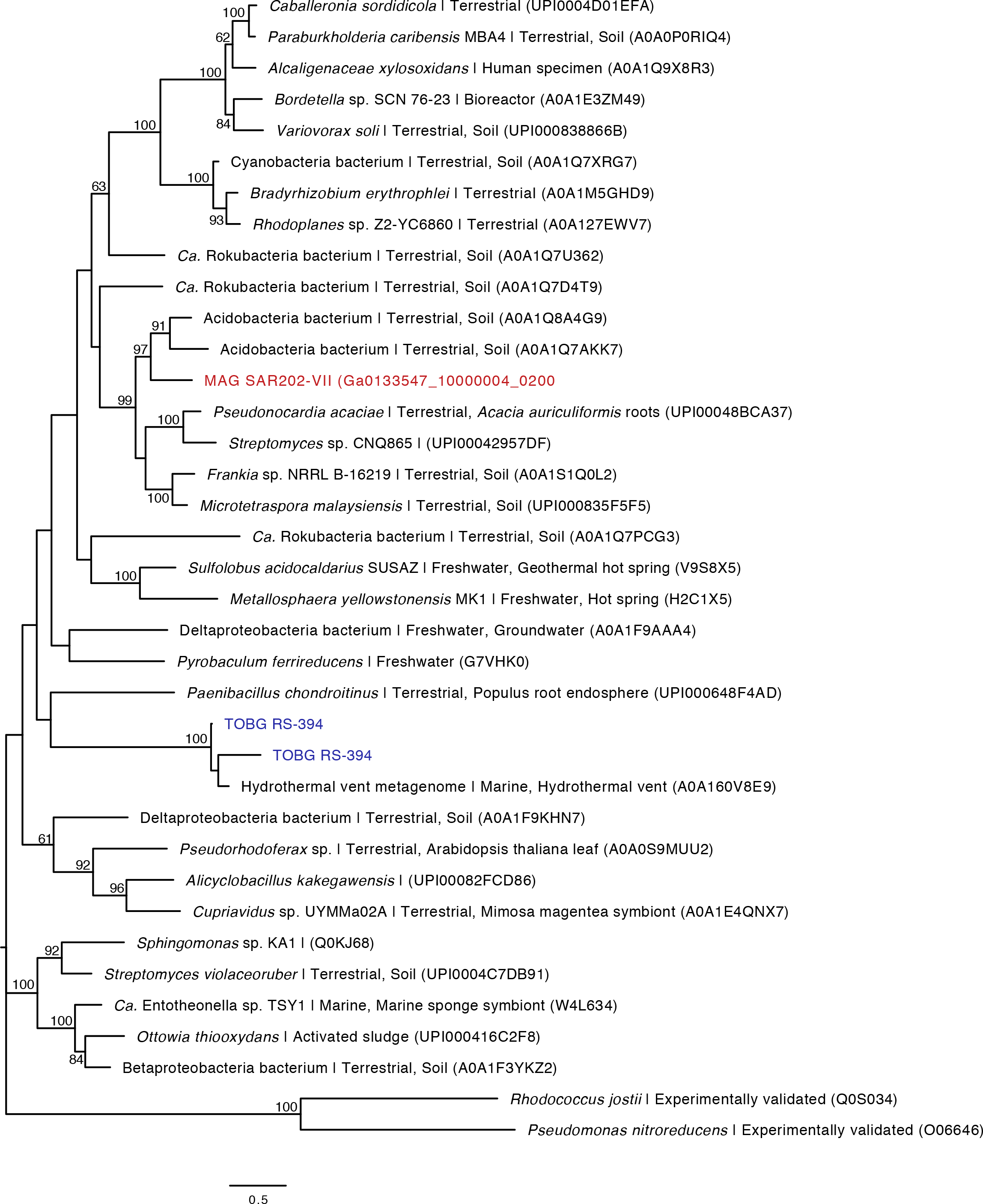
Phylogenetic analysis of a predicted merged LigA/LigB fusion protein sequence identified in SAR202-VII-2. Method and description as described in Supp Figure 2.

**Supplementary Fig 6.**
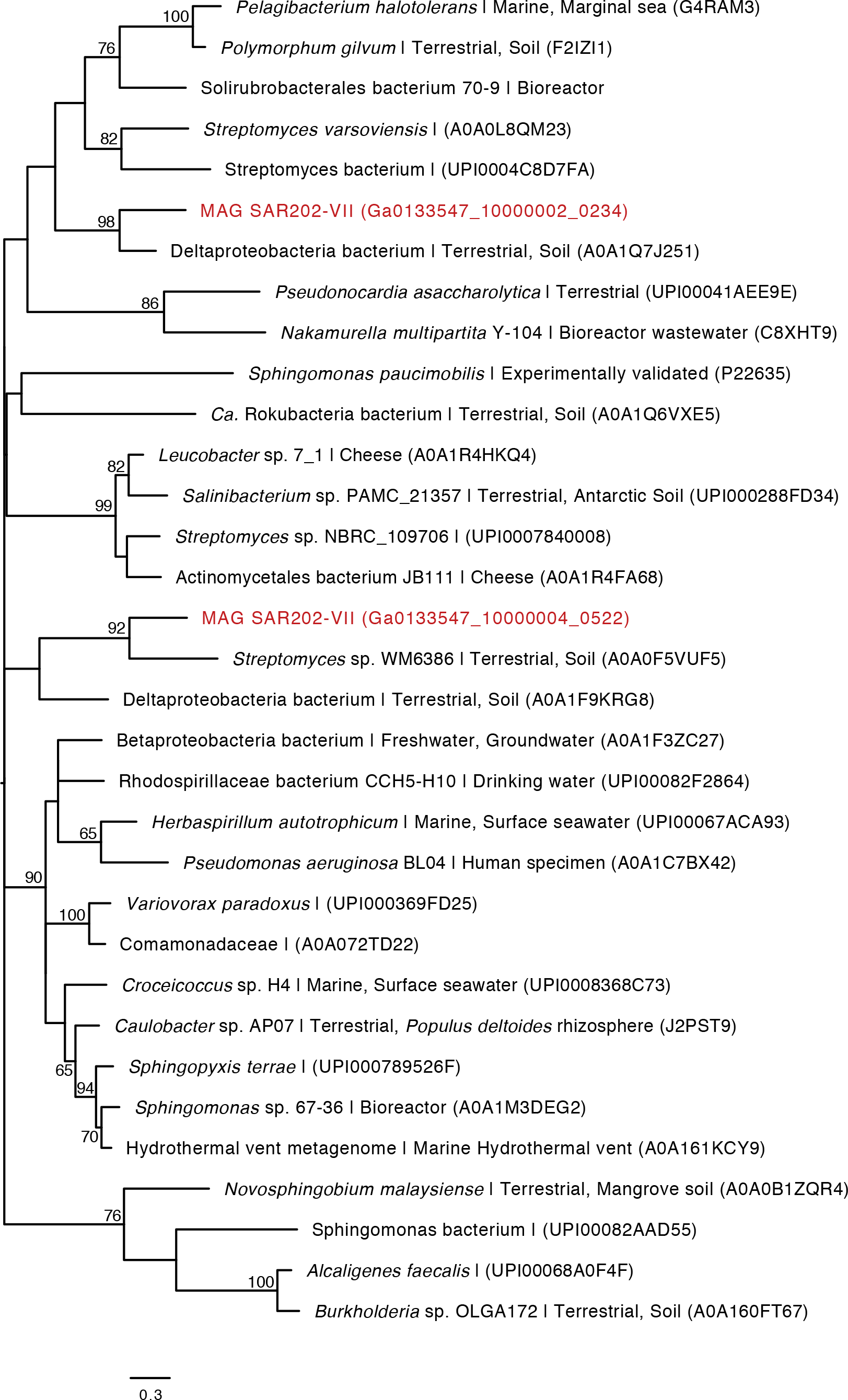
Phylogenetic analysis of predicted LigA dioxygenase (pfam 07746) protein sequences identified in SAR202-VII-2. Method and description as described in Supp Figure 2.

**Supplementary Fig 7.**
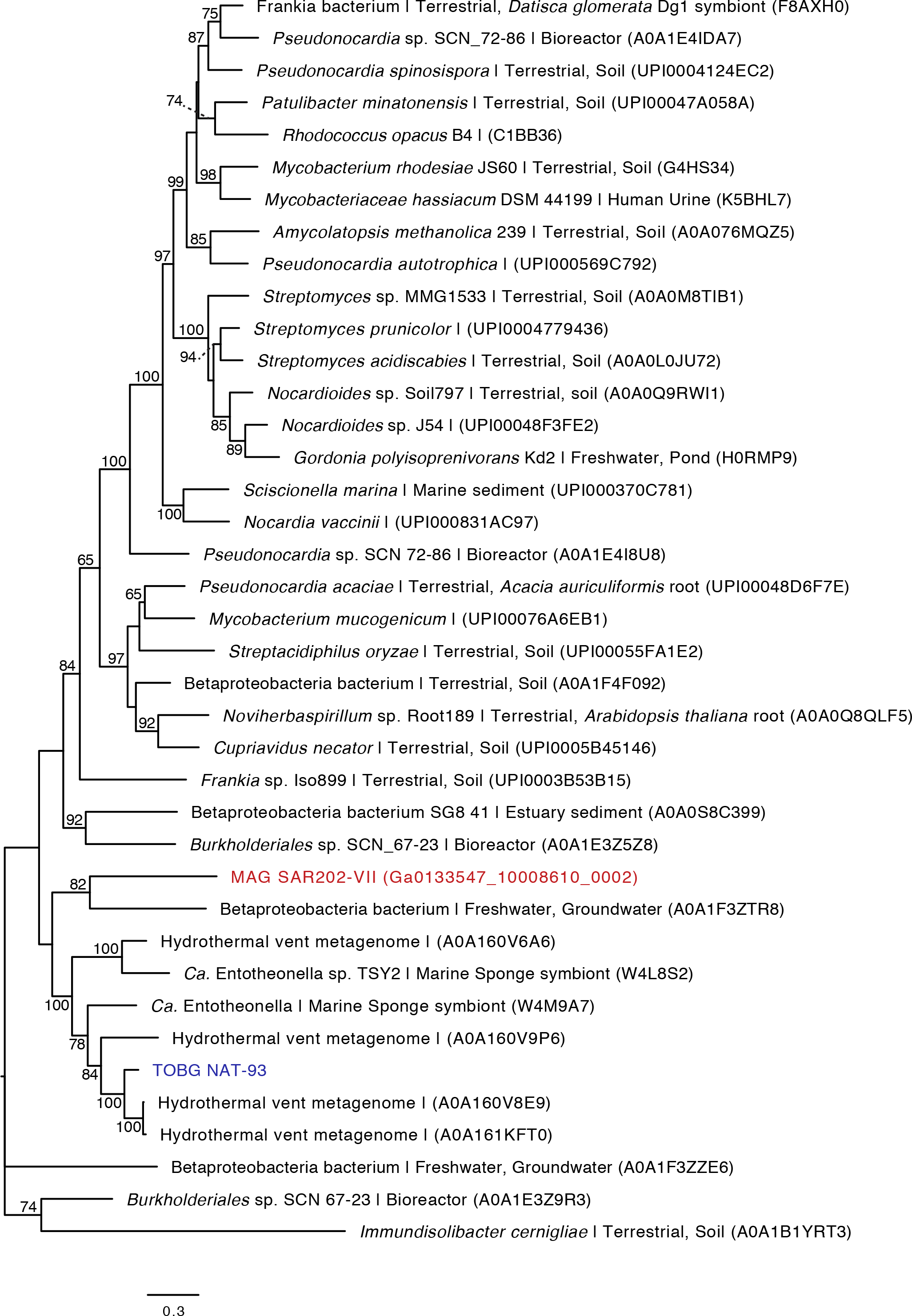
Phylogenetic analysis of a predicted LigB dioxygenase (pfam 02900) protein sequence identified in SAR202-VII-2 (Ga0133547_100086102). Method and description as described in Supp Figure 2.

**Supplementary Fig 8.**
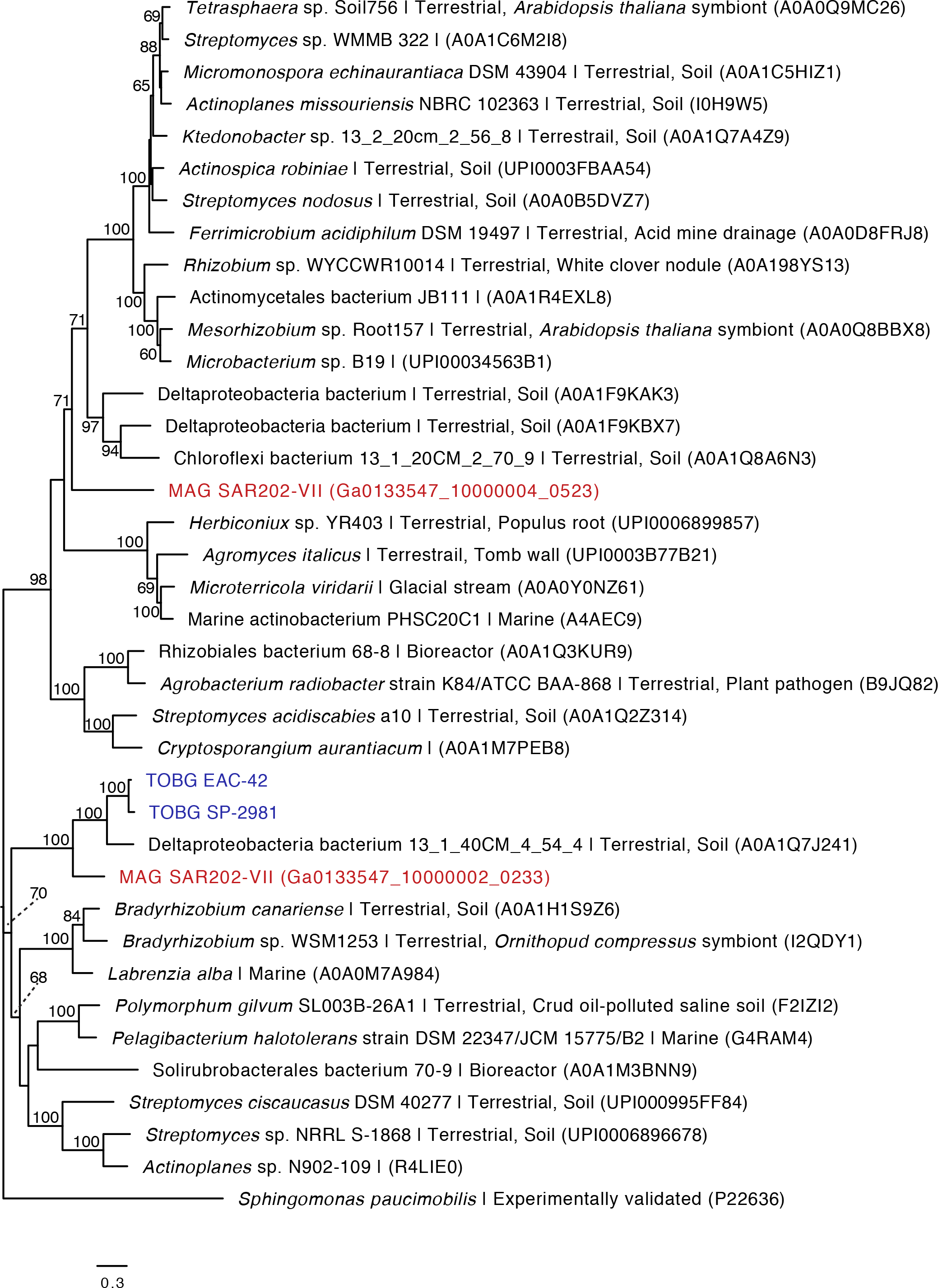
Phylogenetic analysis of predicted beta subunits of the protocatechuate 4,5-dioxygenase (pfam 02900) protein sequences identified in SAR202-VII-2 (Ga0133547_100086102). Method and description as described in Supp Figure 2.

